# PIMMS-Dash: Accessible analysis, interrogation, and visualisation of high-throughput transposon insertion sequencing (TIS) data

**DOI:** 10.1101/2024.04.10.588854

**Authors:** Adam M. Blanchard, Adam Taylor, Andrew Warry, Freya Shephard, Alice Curwen, James A. Leigh, Richard D. Emes, Sharon A. Egan

## Abstract

**Motivation:** Current methods for visualising and interrogating high-throughput transposon insertion mutagenesis sequencing (TIS) data requires a significant time investment in learning bioinformatics, often producing static figures that do not facilitate real time analysis.

**Summary:** We have created an accessible web-based browser tool for visualisation and downstream analysis of high-throughput TIS data. This includes multiple interactive and sortable tables to aid the user to identify genes of interest, enabling the user to gain a greater understanding of the genes contributing to fitness in their experimental work. PIMMS-Dash permits researchers, with any level of bioinformatics knowledge, to interrogate data sets and generate publication quality figures.

**Availability:** PIMMS-Dash is freely available and is accessible online at https://pimms-dashboard-uon.azurewebsites.net

## Introduction

Transposon/insertional mutagenesis is a powerful tool frequently used in microbiology to elucidate genes that contribute to fitness (growth/survival). Disrupting genes by random insertional mutagenesis, provides a resource to achieve a greater understanding of how bacterial genotypes contribute to their observed phenotypes. Approaches involve the construction of mutant libraries containing a high-density of insertions before the library is exposed to the selective condition. Frequency of mutations can then be compared before and after exposure to the selective environment or between selective and non-selective environments. This is achieved by high throughput sequencing, focussed on the genome/insertion junctions, allowing for the quantification of conditionally essential sequenecs within the selective condition.

The main laboratory methods for high-throughput mutagenesis mapping are Insertion Sequencing (INSeq) [1], Transposon Insertion Sequencing (TN-Seq) [2], High-throughput Insertion Tracking by deep Sequencing (HITS) [3], Transposon-Directed Insertion site Sequencing (TraDIS) [4] and Pragmatic Insertion Mutant Mapping System (PIMMS) [5, 6]. Each approach is similar, however there are subtle differences in applicable transposons or insertional elements, mutant library preparation, sequencing approaches and software tools used for bioinformatics analysis. There are numerous bioinformatic tools for analysing the raw sequence data for the multiple different experimental approaches which are summarised elsewhere (Cain et al 2020) but currently there are no tools which analyse the raw data and seamlessly offer statistical analysis and visualisation that can be finely tuned for each dataset.

To complement the currently available tools we have developed PIMMS-Dash, a complementary collection of tools working from a simple browser based interface allowing for quick and easy analysis, statistical evaluation and visualisation of insertional mutation mapping sequence data.

## Implementation

The PIMMS-Dashboard was developed in Python version 3.9 using the IDE Pycharm version 2020.2. The app makes use of following external python packages; numpy (v1.20.1), pandas (v1.2.3), plotly (v4.14.3), dash (v1.19.0). Additionally, DESeq processing is done using R version 4.0.4 and the package DESeq2 (v3.14). Interaction between python and R scripts is handled with the package Rpy2 (v3.4.4). The web app is hosted on the Azure App Service Plan and can be accessed here [https://pimms-dashboard-uon.azurewebsites.net/]. Docker (v20.10.7) was used to containerise the application and push the container image to the Azure container registry.

There are two sets of example data provided on the web app “PIMMS_Example_A”. The respective “control” and “test” files can be selected in the left control panel alongside their “coordinate-gff files” and run through the dashboard. Users can also upload their own csv and gff files which can be generated using PIMMS2 (https://github.com/Streptococcal-Research-Group/PIMMS2) via the upload box, which will make them available to run in the dropdown lists. Any uploaded data is only available to the current browser session and does not become publicly available. It is recommended that the dashboard is explored with the example datasets prior to use with new data to provide an overview of its use. The general dashboard options can be found in the options tab on the left control panel. These include plot configurations for the visualisation tabs and the ability to toggle DESeq processing. We recommend that the default outlier removal in DESeq2 is left enabled.

The PIMMS-Dashboard pre-loaded datasets are from a high-throughput insertional mutagenesis sequencing experiment comparing *Streptococus suis* (P1/7) following growth in laboratory medium (Todd Hewitt broth) and pig serum. New data can be easily uploaded using the drag and drop option on the home screen (Fig 1 A). After loading data the user can work through the tabs to see different analyses and results. The six available tabs allow for the user to visualise the uploaded data table to check its integrity. The NIM Comparison tab allows users to visualise the saturation of insertions across the genomes, between each condition (Fig 1 B). There are also numerous metrics produced from the DESeq2 module which determines the relative fitness of genes between each condition (Fig 1 C). On the Venn tab, users can identify essential genes which are shared or unique to the conditions tested and export a subset list of genes from a chosen intersect (Fig 1 D). The Genome Scatter tab produces an interactive figure where the user can zoom in on regions of the genome to investigate larger areas of essential genes and see if they are represented in both conditions (Fig 1 E). The Replicates tab offers a method of quality control to see if the replicates cluster as would be expected for the two conditions (Fig 1 F). Finally, the GeneViewer tab enables finer scale assessment of the insertions detected in a specific gene, which can be selected in the Data Table tab (Fig 1 G).

**Figure One:**
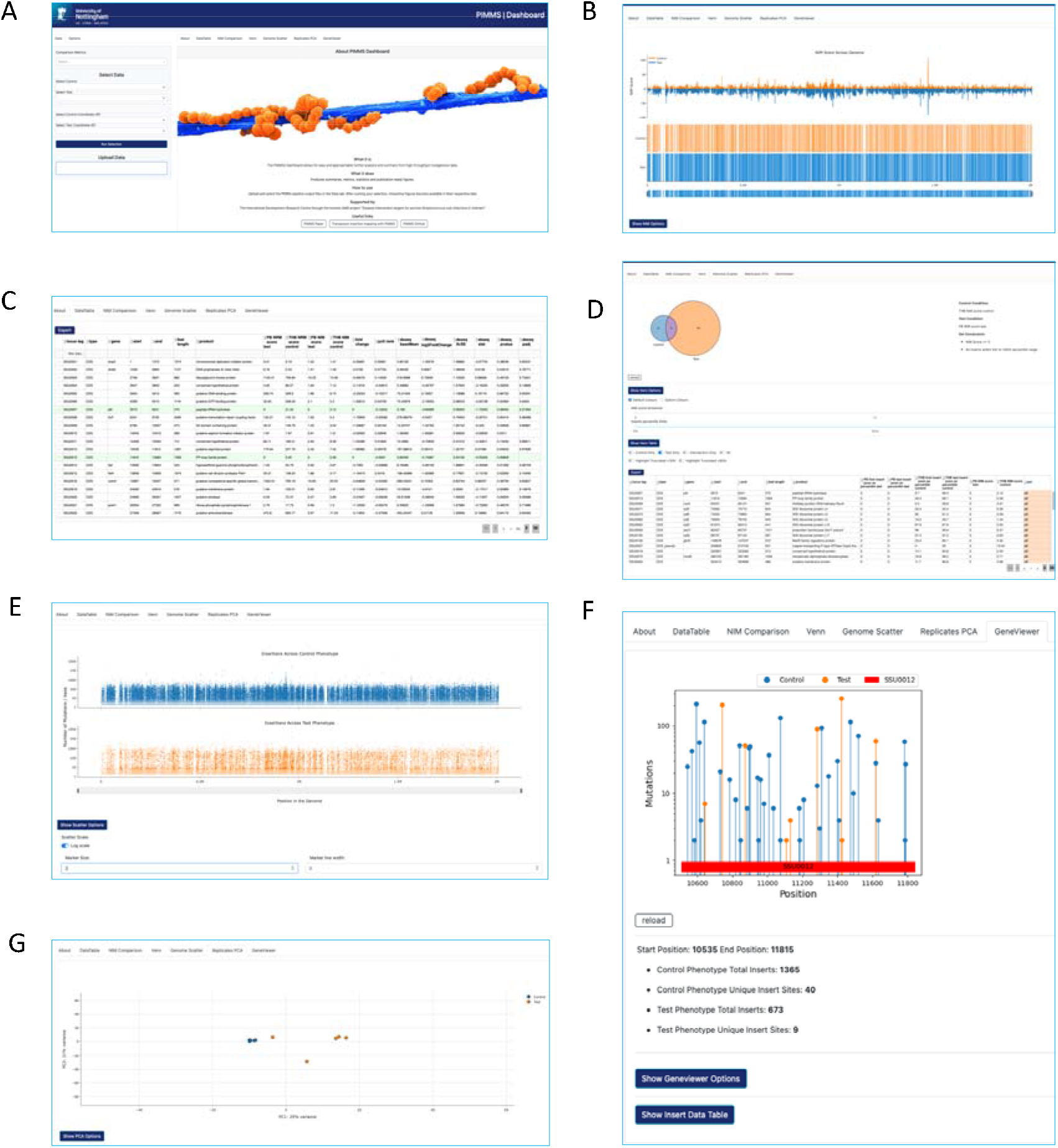
A) Landing page for uploading data and applying settings B) Normalised data to directly compare between test and control mutant libraries and insertional positions. C) Data table with statistical analysis. D) Venn page to identify and subset essential genes. E) Genome Scatter page to visualise regions of genome which lacks any mutations or identify areas of high saturation. F) Gene Viewer page to interrogate the localisation of the insertions at a CDS level. G) Scatter plot to visualise the (dis)similarity of the pools.

## Conclusion

Exploration and analysis of high throughput still remains a daunting task for a bioinformatics novice. PIMMS2 and PIMMS-Dash offers an accessible approach to insertion mapping data analysis with the emphasis on quick and easy statistical analysis and visualisation of results to a publication ready standard.

## Funding Information

This work was supported in part by the International Development Research Centre (IDRC) [Ref: 109056] and the Biotechnology and Biological Sciences Research Council (BBSRC) [Ref: BB/T001933/1 ].

## Notes

### Competing Interest Statement

The authors have declared no competing interest.

